# Htr3a receptors control attenuation of fear responses by modulating the corticolimbic activity and synchronization

**DOI:** 10.64898/2026.03.16.711072

**Authors:** Seblewongel Zewdie Wondimu, Thomas Marissal, Gwenaël Birot, Karl Schaller, Alexandre Dayer, Charles Quairiaux

**Author notes:** contributed equally (Alexandre Dayer, Charles Quairiaux).

## Abstract

The fear circuit orchestrates defensive responses to environmental threats and is essential for survival. Dysregulation of this system is thought to contribute to the pathophysiology of several psychiatric disorders. Within this fear circuit, the corticolimbic network, particularly the amygdala and the medial prefrontal cortex (mPFC), is strongly modulated by serotonin. Previous studies have shown that *Htr3a* knockout (*Htr3a-KO*) mice exhibit deficits in the extinction of cued fear memory; however, the circuit level mechanisms underlying these impairments remain unknown. Here, we investigated this question by recording local field potentials evoked by auditory conditioned stimuli (CS) in the prelimbic (PrL), infralimbic (IL), and basolateral amygdala (BLA) of head-fixed wild-type (WT) and *Htr3a-KO* mice prior to fear conditioning and during fear memory retrieval. Behaviorally, *Htr3a-KO* mice displayed a delayed attenuation of fear-induced freezing during cued fear memory retrieval, whereas WT mice showed a rapid attenuation in freezing. Electrophysiologically, *Htr3a-KO* mice exhibited reduced fear-evoked theta power in the PrL, IL, and BLA, along with diminished mPFC-BLA theta synchrony. Moreover, theta-phase modulation of gamma oscillations within the BLA, which has been shown to increase during fear states, was perturbed in the absence of Htr3a signaling. Together, these findings indicate that Htr3a is critical for maintaining proper oscillatory dynamics within the mPFC-BLA circuit and for supporting effective attenuation of learned fear.

**Highlights:**

- Attenuation of fear responses during fear memory retrieval sessions is protracted in *Htr3a* knock-out mice
- The fear-induced theta response in the medial prefrontal cortex and the basolateral amygdala is less powerful in the Htr3a knock-out mice than in wild-type
- *Htr3a* knock-out mice show a deficit in fear-induced synchronization as well as in theta modulation of gamma power in the cortico-limbic network
- These results suggest that malfunction of the Htr3a receptor cause alterations in fear network circuit mechanisms that might be linked to deficits in fear responses attenuation

## Introduction

The brain can associate sensory stimuli with threats, forming long-lasting fear memories that enable us to respond appropriately to danger and are essential for survival. Dysfunctions in the brain circuits involved in fear processing can lead to excessive, inappropriate, or persistent fear reactions, contributing to mental disorders such as anxiety disorders, posttraumatic stress disorders (PTSD), depression, and phobias (Jovanovic and Ressler, 2010; Bukalo et al., 2014; Gasparyan et al., 2022). Therefore, studying the mechanisms underlying fear modulation is highly important from a clinical perspective.

The classical fear conditioning paradigm, in which an aversive unconditioned stimulus (US) is paired with a conditioned stimulus (CS) to elicit a behavioral response, is one of the most common tools for studying the neuronal substrates of associative fear learning (Tovote et al., 2015). This paradigm has enabled the identification of brain circuits involved in fear memory acquisition, expression and extinction of fear memories. Extinction refers to the attenuation of the conditioned fear response when the CS is repeatedly presented without the US (Myers and Davis, 2007; Tovote et al., 2015; Ferrara et al., 2023). However, the mechanisms that support this long-lasting reduction in fear responses are complex and remain the focus of intensive research aimed at improving behavioral interventions for anxiety disorders (Craske et al., 2014; McGlade et al., 2023).

The neuronal circuits involved in fear memory acquisition, retrieval, and extinction are distributed across corticolimbic structures. Among these, the basolateral amygdala (BLA) and the prelimbic (PrL) and infralimbic (IL) regions of the medial prefrontal cortex (mPFC) are key regulators of fear memory (Maroun, 2013; Likhtik et al., 2014; Stujenske et al., 2014; Karalis et al., 2016; Whittle et al., 2021). Numerous studies have shown that synchronous oscillations within the BLA-mPFC circuits, particularly in the theta-range, correlate with experience-dependent fear behavior and modulate its expression (Lesting et al., 2011, 2013; Likhtik et al., 2014; Stujenske et al., 2014; Karalis et al., 2016; Ozawa et al., 2020).

These regions contain dense serotonergic projections. Serotonin (5-HT) levels in the amygdala have been shown to increase during both cued and contextual fear conditioning, modulating activity within these circuits (Bauer, 2015). This modulatory effect is mediated by various 5-HT receptors, including the ionotropic Htr3a receptor that is particularly expressed in interneurons of the cortex and amygdala (Mascagni and McDonald, 2007; Rudy et al., 2011). Interestingly, Htr3a receptors are necessary for the extinction, but not the acquisition and retention, of contextual or tone-cued fear (Kondo et al., 2014).

However, the circuit mechanisms by which Htr3a receptors control the extinction of fear are unknown. Here, we focus on the early mechanisms underlying the attenuation of conditioned fear responses. To this end, we performed auditory cued fear conditioning and *in vivo* multiple-site intra-cerebral electrophysiological recordings to investigate local field potential (LFP) dynamics and activity synchronization in the BLA and mPFC of Htr3a-deficient (*Htr3a-KO*) mice during retrieval. We observed that the attenuation of fear responses across repeated CS presentations was delayed in freely moving *Htr3a-KO* mice compared to wild-type animals. LFP recordings performed on head-fixed mice show that this delayed attenuation of fear responses (observed using pupillometry) is associated with alterations in neuronal responses to the CS, including a decrease in theta amplitudes in BLA and mPFC and reduced mPFC-BLA theta phase locking in Htr3a-deficient animals. Therefore, we propose that Htr3a receptors control fear attenuation by modulating the activity and synchronization of the corticolimbic fear network.

## Material and Methods

Twenty male C57BL/6j adult WT and Htr3a-KO mice (12-20 weeks) were included. Experiments were performed in accordance with the Geneva and Switzerland animal care committee’s regulations.

### Head-fixed surgeries

A head-fixed protocol was used to minimize movement artifacts during multiple intracerebral recordings. Mice were anesthetized with a 60 ml intra-peritoneal (IP) injection containing 0.23 mg/ml Domitor (medetomidine hydrochloride), 3 mg/ml Dormicum (midazolam), and 7.7 µg/ml Fentanyl Sintetica. Mice were then placed on a homeothermic blanket (Harvard Apparatus) and secured on a stereotaxic apparatus (Narishige). 2.5 mg/ml of a preoperative analgesic was administered subcutaneously (100 µl Carbostesin; Bupivacaine hydro-chloridum). A head-fixation bar was attached to the skull using dental cement and a reference electrode was implanted in the caudal end of the right parietal bone at −4.2 mm anteroposterior (AP) from bregma and −0.5 mm mediolateral (ML) from midline. Craniectomy was performed using a drill above Prl and IL in mPFC (coordinates: 1.7 mm AP and 0.5 mm ML) as well as BLA (−1.7 mm AP and 3 mm ML) to enable the electrophysiological recordings. The exposed skull was protected using Kwik Cast sealant and dental acrylic (Paladur). At the end of the procedure, mice received a 250 ml subcutaneous injection of a waking solution containing 97.8 µg/ml Anexate (imidazobenzodiazepine derivative), 300 µg/ml Alzane (Atipamezole hydrochloride), 5.8 µg/ml Naloxon (Naloxoni hydrochloridum). Postoperative analgesics and antibiotics were provided (Dafalgan Paracitamol [250 mg/250 ml], Algifor Junior [2.5 ml/250 ml], Nopil [5 ml/250 ml]) in drinking water. Mice were allowed to recover from surgery for 4-7 days.

### Habituation and cued fear memory retrieval in head-fixed mice

Mice were handled daily by the experimenter for at least 3 days prior to the start of the protocol. From Day 1 to Day 3 (D1–D3), mice were habituated to the head-fixation apparatus for 30 minutes, 1 hour, and 2 hours, respectively. During habituation, mice were exposed to 14 auditory conditioned stimuli (CS: trains of 30 white noise 7.5 kHz and 70 dB pips of 50 ms in duration, delivered at 1 Hz over 30 seconds) with a pseudorandomized inter-stimulus interval (ISI) (Likhtik et al., 2014). Pupil size and local field potential (LFP) recordings were conducted on D4 and D6, corresponding to the habituation and cued fear memory retrieval sessions, respectively, as detailed below.

### Fear conditioning

On D5, mice underwent fear conditioning in a sound-attenuating conditioning chamber (17 × 17 × 25 cm; Ugo Basile, 46003), equipped with a grid floor of stainless-steel bars for shock delivery. Mice were presented with the conditioned stimulus —30-second train of 50 ms/70 dB/7.5 kHz pips delivered at 1 Hz as for the habituation day— paired with an unconditioned stimulus (US)—a 0.6 mA foot shock delivered to the paws for 2 seconds. Mice were allowed to explore the conditioning chamber for 120 seconds before the protocol began. They then received 5 CS presentations, each ending simultaneously with the foot shock. Each CS–US pair was followed by a pseudorandomized inter-stimulus interval ranging from 60 to 90 seconds. Following the final CS–US pairing, mice were returned to their home cage after a 30-second delay. The conditioning chamber was cleaned with 70% ethanol and 1% acetic acid before and after each session to minimize olfactory cues. Sessions were recorded with an overhead camera and the time spent by each mouse freezing was calculated using AnyMaze behavior tracking software (Stoelting). Freezing behavior was defined as the absence of any movement for at least 2 seconds.

### Fear memory retrieval in freely moving mice

On D7, cued and contextual fear memory recall was assessed in freely moving mice. To assess cued fear memory, mice were placed in a novel context chamber (57L x 47W x 54H cm) and were re-exposed to the 14 CS with pseudorandomized ISI. Subsequently, mice were returned to the original conditioning chamber (initial context) in the absence of the CS to assess contextual fear memory recall.

### Pupillometry in head-fixed mice

Pupil size was monitored in head-fixed mice on D4 (habituation) and D6 (cued fear memory retrieval) with a digital USB camera (CMOS camera, 1280 × 1024, Monochrome, ThorCam, Germany) with a 1.00X Bi-Telecentric C-mount lens (Thorcam, Germany). Images were acquired and saved at 8 frames/s (240×376 pixels 8-bit greyscale images) using ThorCam video acquisition system. Infrared (IR) light (940nm LED, Everlight Electronics, 3 mW, 40 mA, 1.2 V) was illuminated on the eye to obtain high-contrast images of the pupil. IR was switched on and off by a transistor-transistor-logic (TTL) from the AnyMaze software at the start and end of each session to mark the first and last frames of pupil recording, respectively.

Pupil images were processed using Matlab image processing toolbox and custom-written scripts. To extract the pupil diameter, single frame images were first centered on, cropped around the pupil, and processed using Top-hat and Gaussian filters. Subsequently, pupil radius was automatically detected within a given range selected a priori based on the minimum and maximum pupil size recorded for each mouse. Post-hoc analyses were used to exclude incorrect identification of the pupil based on the X and Y positions of the circle identified as a pupil. Missing pupil data were then interpolated and subsequently verified for accuracy. The pupil traces were then aligned to CS triggers based on the first and last frames marked by the switching on and off of the IR light. To obtain CS-induced pupil increase, pupil traces during the 30 s CS presentation were normalized as a percent increase from the 10 s period preceding it.

### Electrophysiological recordings

Awake head-fixed intra-cerebral LFP recordings were acquired using a Digital LynxSX amplifier (Neuralynx), Neuronexus A16 probes, and a unitary gain head-stage preamplifier (HS-16, 100µm between contacts, Neuralynx). Linear electrode probes were inserted in the mPFC (+1.7 mm AP, −0.5 mm ML, lowermost electrode at 2 mm depth) to target the prelimbic (PrL) and infralimbic (IL) regions and in BLA (−1.7 AP, 3 mm ML, lowermost electrode at 4 mm depth). Field potential signals were acquired at 16 kHz with a low-pass filter of 8 kHz. Before recordings, animals were briefly anesthetized with isoflurane (1%) to allow for the precise positioning of the electrodes. Recordings were started 30 min after the animal fully woke up to allow for the effects of anesthesia to wear off.

### Local field potential analyses

#### Electrodes selection

Based on the mouse brain atlas (Paxinos and Franklin, 2019), the PrL and IL electrodes were selected from the mPFC probe at 2000 µm and 1600 µm depth and the BLA electrode was selected from the BLA probe at a depth of 3700 µm. If one of the channels was broken, we selected the channel immediately above or below, i.e. at −/+ 100 µm. The signal from each electrode was then re-referenced to the uppermost electrode of the probe. Data were down sampled to 1000Hz, band pass filtered (1-100 Hz) and analyzed using custom-written script based on Matlab (R2017A, MathWorks) signal processing toolbox along with functions in Chronux open-source software package. LFPs were averaged across trials in a window centered on the pip-onset and the square-root of the voltage was calculated to create a pip-evoked potential.

#### Spectral analyses

Power spectra and spectrogram were calculated using a multitaper method with a moving window of 375ms and 5ms overlap, and time-bandwidth product of 1.5 (mtspectrumc_mod). Theta power was defined as the 4-12 Hz frequency band. Coherence was calculated with a multitaper method using a moving window of 250ms with 5ms overlap, a time-bandwidth product of 1.5 and FFT padding factor of 5 (cohgramc). Coherence comprises both power correlations and phase consistency (i.e., phase coherence) within the same frequency band. As phase consistency can vary independently of power correlations, it was analyzed separately.

#### Theta phase synchrony

Theta phase synchrony between PrL, IL, and BLA was detected with an open-source script using a filter order 50. Instantaneous phase of every signal at theta frequency was extracted through Hilbert transform and for each latency a measure of phase-locking was computed at each frequency (phase-locking value, PLV) (Lachaux et al., 1999).

#### Phase-amplitude coupling

Instantaneous values for theta phase and gamma amplitude were obtained using Hilbert transform on band pass filtered LFP data. The strength of the modulation of gamma amplitude by theta phase was measured by first normalizing to the mean gamma power and then computing the fractional modulation of gamma amplitude by theta phase. Modulation index (MI) was derived from the Kullbach-Leibner divergence and was used to measure the amount of phase-amplitude coupling between theta and gamma signals (Tort et al., 2010).

### Statistical analyses

Statistical analyses were performed using GraphPad Prism software. The normality of data distributions was assessed using the Shapiro-Wilk method. For fear acquisition and retrieval in freely moving mice, freezing responses were analyzed using two⍰way repeated⍰measures ANOVA, with the repetition of CS trains, including baseline, as the within⍰subjects factor (CS trains) and the mouse group, WT vs Htr3a⍰KO, as the between⍰subjects factor (Genotype). Because comparisons between baseline and subsequent CS trains were planned a priori, paired t⍰tests were performed following a significant main effect of CS trains, without additional correction for multiple comparisons. Paired t⍰tests were used to compare CS⍰induced freezing responses averaged across all CS trains in Figures⍰1E and 1F, except for the Htr3a⍰KO group in Figure⍰1I, for which a Wilcoxon signed⍰rank test was used due to violation of normality assumptions. For the head⍰fixed experiments, the effects of Day (habituation vs retrieval) and CS trains were analyzed using two⍰way ANOVA. Post⍰hoc comparisons between genotypes across CS presentations or across regions were corrected using the Bonferroni method. In addition, for the first CS presentation block, the effects of Day across regions or the effects of genotype across regions were assessed using two⍰way ANOVA followed by Bonferroni⍰corrected multiple comparisons.

## Results

### *Htr3a-KO* mice show reduced attenuation of CS-induced freezing response

We first investigated the contribution of Htr3a receptors to distinct aspects of fear processing by performing an auditory-cued fear conditioning protocol in freely moving Htr3a knockout (KO) and wild-type (WT) mice. Both *Htr3a-KO* and WT mice exhibited a progressive increase in freezing behavior during CS presentations (i.e., each CS consisted of 30 white noise pips of 50 ms durations delivered at 1Hz) compared to the 2-minute baseline period (Fig. 1A, B), indicating that fear memory acquisition does not depend on Htr3a. Next, we assessed whether *Htr3a-KO* mice differed from WT mice in fear memory retrieval in a novel context. Regardless of genotype, mice exhibited a significant CS-induced freezing response: in both strains, average CS-induced freezing across all presentations was significantly higher than baseline (Fig. 1E, F).

**Figure 1:**
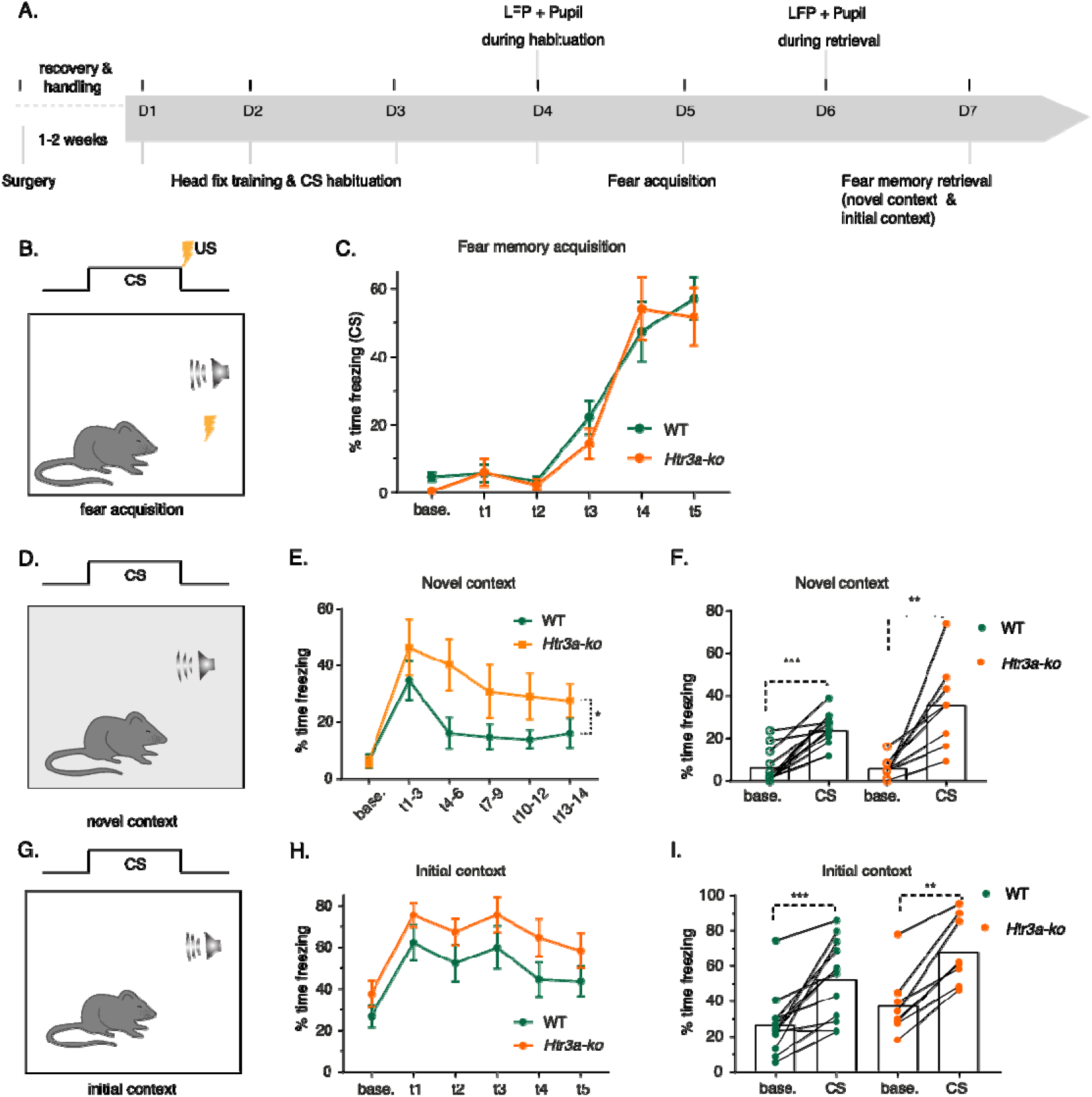
Fear acquisition and retrieval in freely moving WT and Htr3a-KO mice. A) Experimental timeline. B) Fear acquisition setup. C) Percentage of time freezing across Conditional Stimulus (CS) presentations on fear memory acquisition day in WT (n = 12 mice) and Htr3a-KO (n = 8 mice) compared to baseline (main effect of CS p < 0.001, two-way repeated measures ANOVA and post-hoc comparisons; WT base vs t3: p = 0.01, base vs t4 and base vs t5: p < 0.001; Htr3a-KO base vs t3: p = 0.213, base vs t4 and base vs t5: p < 0.001). D) Fear memory retrieval in a novel context set up. E) CS-induced freezing response during fear memory retrieval in a novel context in WT and Htr3a-KO animals across 14 CS presentations as compared to baseline. Main effect of genotype *p = 0.037 and main effect of CS p < 0.001 in two-way repeated measures ANOVA. Paired t-tests comparing baseline to CS train blocks showed significantly higher freezing for the first CS block in WT mice, whereas subsequent blocks did not differ significantly from baseline (base vs t1-3: p = 0.002, base vs t4-6: p = 0.068, base vs t7-9: 0.142, base vs t10-12: p = 0.051, base vs t13-14: p = 0.053). In Htr3a-KO mice, freezing remained significantly higher than baseline for all CS blocks (base vs t1-3: p = 0.006, base vs t4-6: p = 0.005, base vs t7-9: p = 0.036, base vs t10-12: p = 0.030, base vs t13-14: p = 0.007). F) CS-induced freezing response averaged across presentations in WT and Htr3a-KO compared to baseline during fear memory retrieval in novel context. **p = 0.006, ***p < 0.001, Paired t-test. G) Fear memory retrieval in the initial context setup. H) Freezing response during fear memory recall in the initial context in WT and Htr3a-KO mice. Main effect of CS p < 0.001 and main effect of genotype p = 0.113 in repeated measures ANOVA, followed post-hoc comparisons (WT: base vs t1: p < 0.001, base vs t2: p = 0.004, base vs t3: p < 0.001,base vs t4: p = 0.094, base vs t5: p = 0.126; Htr3a-KO: base vs t1: p < 0.001, base vs t2: p = 0.008, base vs t3: p < 0.001, base vs t4: p = 0.019, base vs t5: p = 0.123). I) CS-induced freezing response averaged across presentations compared to baseline in WT and Htr3a-KO in the initial context. Paired t-test and Wilcoxon test. **p < 0.01, ***p < 0.001. All graphs mean ± SEM.

Figure 1EF show the percentage of time spent freezing, averaged either across all CS presentations (Fig. 1F) or across blocks of presentations (Fig. 1E), during baseline (base.) and retrieval in a novel context. Both WT and *Htr3a-KO* mice displayed increased freezing during retrieval. A repeated-measures ANOVA revealed a main effect of CS trains and a significant effect of genotype, consistent with a slower attenuation of CS-induced freezing in *Htr3a-KO* mice. In both groups, freezing decreased after the first blocks of CS presentations; however this reduction was more pronounced in WT mice, whereas freezing remained significantly higher than baseline across all CS blocks in *Htr3a-KO* mice (Fig. 1E). This genotype-dependent difference in freezing response in a novel context suggests that Htr3a contributes to the attenuation of CS-induced freezing during fear retrieval. When mice were placed back in the initial fear-acquisition context, both WT and *Htr3a-KO* groups continued to display CS-induced freezing response during subsequent CS presentations (Fig. 1H, I), indicating intact contextual fear memory in *Htr3a-KO*. Taken together, and consistent with previous work (Kondo et al., 2014), our results indicate that Htr3a is essential for the attenuation of cue-induced fear responses.

### mPFC and BLA LFP responses to the CS during retrieval are impaired in *Htr3a-KO* mice

LFP responses to the auditory cue were monitored during head-fixed recordings in the corticolimbic fear circuit at D4 and D6, i.e., 24 h before and after fear conditioning, in both WT and *Htr3a-KO* groups. We measured CS-induced pupil dilations as a behavioral read-out to ensure that CS-fear responses could be induced in this distinct experimental setting (see Supplementary figure S1). Figures 2CD illustrate the pip-induced LFP across successive CS presentations in the PrL, IL and BLA regions. WT mice showed an increased pip-evoked activity in all 3 regions during the retrieval day compared to the habituation day. This increase is more prominent during the presentations of the first 3 CS trains (t1-t3; Fig. 2C, 2E). However, *Htr3a-KO* mice did not show any significant main effect between habituation and retrieval days (Fig. 2D, 2E). While no significant differences in pip-induced LFPs were observed between WT and *Htr3a-KO* mice during habituation, *Htr3a-KO* mice exhibited weaker pip-evoked activities than WT during retrieval, this difference being more robust in the BLA (Fig. 2F). Thus, fear conditioning induces an increase in CS evoked activity in the mPFC and BLA regions in WT mice, which progressively attenuates across CS presentations. In *Htr3a-KO* mice, however, no significant increase in CS-evoked responses was observed after fear conditioning, and the attenuation dynamics were absent.

**Figure 2.**
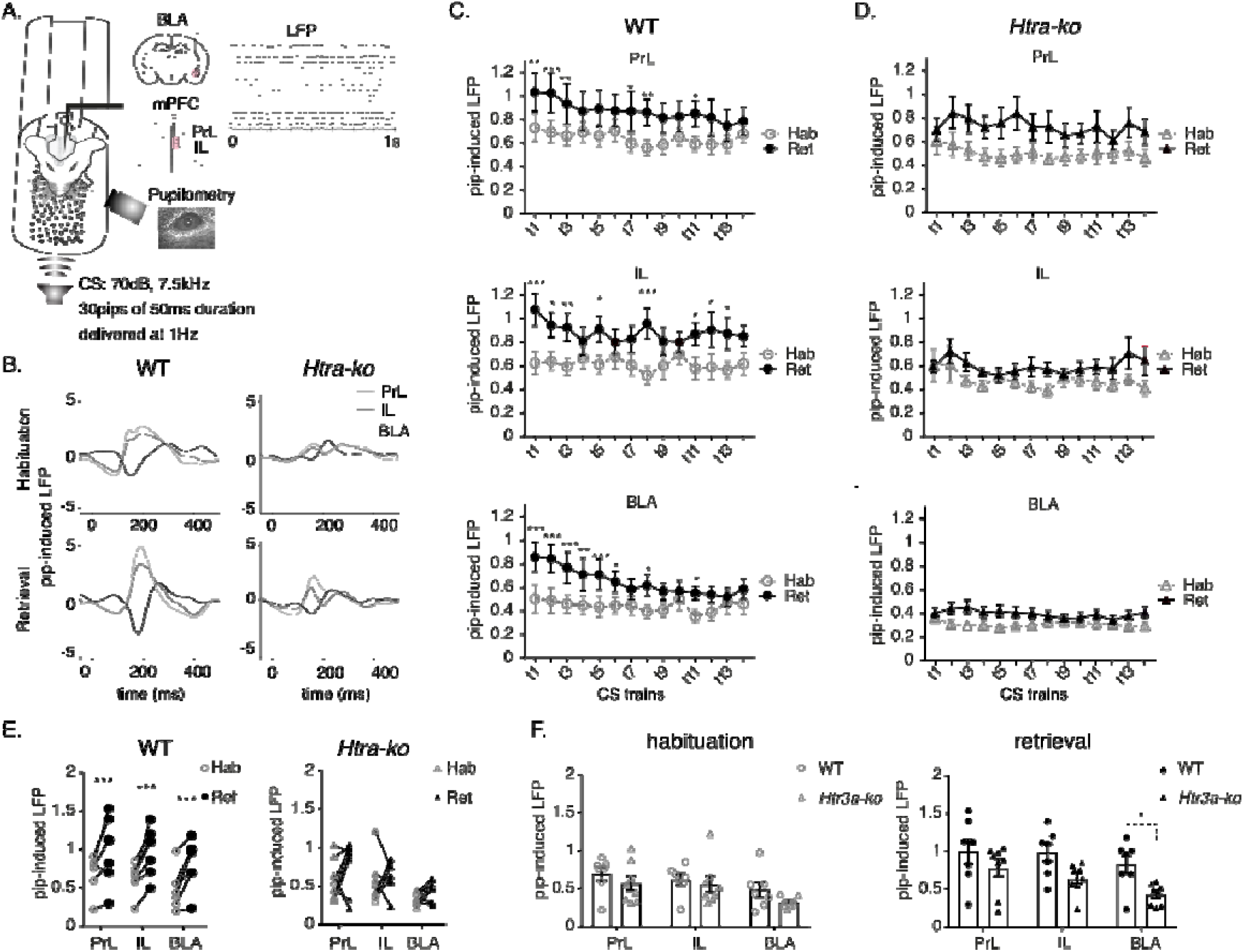
Fear conditioning increases pip-induced local field potential (LFP) in control but not in Htr3a-KO during retrieval. A) Schema of the experimental setup for intracerebral recordings in head-fixed mice in the prelimbic cortex (PrL), infralimbic cortex (IL) and basolateral amygdala (BLA). B) Representative examples (1 mouse per condition) of pip-evoked potentials (1-100 Hz), z-normalized to 500ms pre-pip period, in WT and Htr3a-KO mice. Mean of all CS trains (420 pips, solid lines) ± SEM. C) Average pip-evoked LFP ± SEM across the 30 pips of each CS train in WT mice (n = 8; LFP data of 4/12 WT mice were excluded due to poor signal quality) in the PrL, IL, and BLA during habituation and retrieval days. Data are shown across the 14 CS trains (labelled t1 to t14) and were z-normalized to the 500 ms pre-pip baseline. In the PrL, there was a main effect of Days (p = 0.049) and of CS train (p = 0.002). In the IL, there was a main effect of days (p = 0.04), whereas the effect of CS train was not significant (p = 0.77). In the BLA, there was a main effect of days (p = 0.009) and of CS train (p < 0.001). Two-way ANOVA followed by Bonferroni post-hoc tests (*p < 0.05, **p < 0.01, ***p < 0.001). D) Pip-evoked responses in the Htr3a-KO mice (n = 8), no main effects were detected. E) Averaged LFP response of the first 3 CS during habituation and retrieval days, z-normalized to the 500ms pre-pip-onset in WT (main effect of Days: p =0.004, Bonferroni multiple comparison for PrL, IL and BLA: all p < 0.001) and Htr3a-KO (main effect of Days: p = 0.201). F) LFP responses of the first 3 CS in all mice during habituation (main effect of genotype: p = 0.275) and during retrieval (main effect of genotype: p = 0.038, Bonferroni multiple comparison for PrL: p = 0.489, IL: p = 0.097, BLA: *p = 0.043).

### Theta power fails to increase in the fear network of *Htr3a*-KO mice during fear memory retrieval

Theta power is known to increase in the fear network during fear retrieval (Karalis et al., 2016; Taub et al., 2018). We therefore examined whether theta activity (4-12 Hz) increased in the BLA and both subregions of the mPFC during CS presentations in the presence or absence of functional Htr3a receptors. Time-frequency plots confirmed the LFP observations, showing prominent pip-evoked activities in both WT and *Htr3a-KO* mice (Fig. 3A) during habituation and retrieval days around the theta range and between 15–30 Hz, the latter likely reflecting theta harmonics rather than distinct activity (Sheremet et al., 2016). In WT mice—but not in KO mice—we observed that evoked theta power was greater on the retrieval day than during habituation, with a significant effect of recording day in all three regions (Fig.⍰3C). This effect in WT animals was strongest during the first three CS trains, although the repetition effect (CS train) reached statistical significance only in the BLA (p⍰<⍰0.001). An analysis focusing on the average response during the first three CS trains revealed a significant fear-conditioning effect in all regions for WT mice, but not for *Htr3a-KO* mice (Fig.⍰3E). Accordingly, while no genotype differences were detected during habituation, a significant genotype effect emerged during retrieval: *Htr3a-KO* mice exhibited lower pip-induced theta power than WT mice, reaching statistical significance in the BLA (Fig.⍰3F).

**Figure 3.**
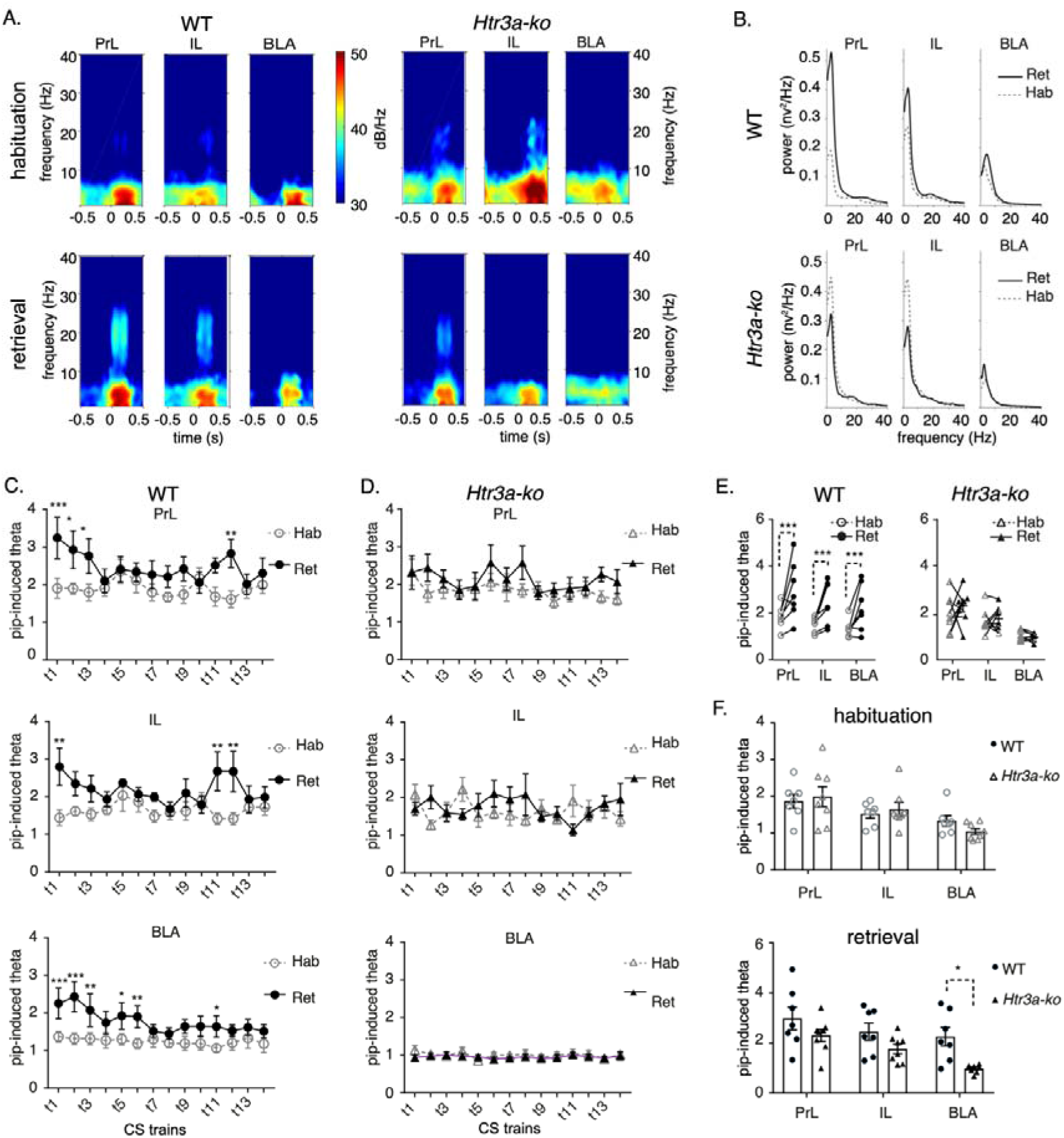
Theta power fails to increase during retrieval in Htr3a-KO mice. A) Representative examples of spectrograms showing pip-induced low frequency activity in Prelimbic (PrL, left), Infralimbic (IL, centre) cortices and Basolateral Amygdala (BLA, right) during presentation of the first train on habituation (top) and retrieval (bottom) days in WT and Htr3a-KO. B) Representative examples of power spectral density of 500 ms post pip-onset averaged across the first 3 CS trains in a WT or a Htr3a-KO mouse. C) Theta (4-12 Hz) power increase from pre-pip period across the 14 trains (t1 to t14) in PrL (effect of days: p = 0.04), IL (main effect of days: p = 0.003) and BLA (main effect of Days: p = 0.018). D) Theta (4-12 Hz) power increase from pre pip-onset period across 14 trains in PrL, IL and BLA in Htr3a-KO mice, no main effects were detected. E) Comparison between average pip-induced theta power of the first 3 CS trains during the habituation and retrieval days in WT (effect of Days: p = 0.017) and Htr3a-KO mice (effect of Days: p = 0.58). F) Comparison between average pip-induced theta power of the first 3 CS trains in WT as compared to Htr3a-KO mice in the habituation (effect of genotype: p = 0.946) and retrieval (effect of genotype: p = 0.02) days. Theta power was significantly lower in the BLA of Htr3a-KO as compared to WT during the retrieval day (p = 0.012). All analyses used two-way ANOVA with Bonferroni multiple comparisons. Graphs are mean ± SEM. *p < 0.05, **p < 0.01, ***p < 0.001.

### Reduced mPFC-BLA theta phase locking during fear memory retrieval in *Htr3a-KO* mice

Given that theta power was enhanced during retrieval in the mPFC regions and the BLA of WT but not *Htr3a-KO* mice, we further investigated theta phase locking between the mPFC and BLA regions. Figure 4A shows post-pip theta phase-locking values (PLV) during the 500 ms post-pip period, normalized to the pre-stimulus baseline, for habituation and retrieval days. These data reveal an increase in phase locking, particularly pronounced on the retrieval day in the IL-BLA and PrL-BLA networks of WT mice.

**Figure 4.**
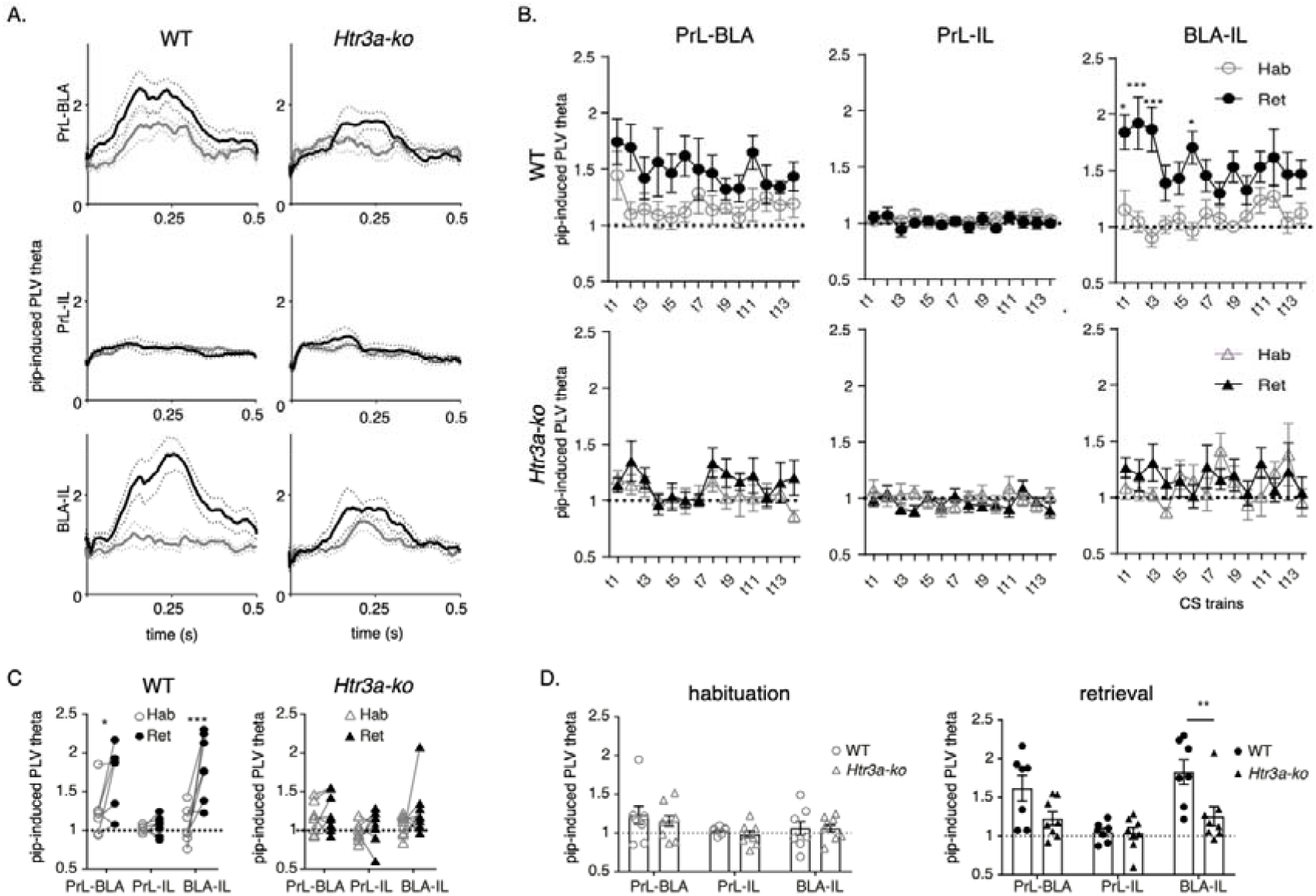
mPFC-BLA theta phase-locking during fear memory retrieval is impaired in Htr3a-KO mice. A) Cross regional post-pip (500 ms) phase locking of theta (4-12 Hz) in WT (n = 8) and Htr3a-KO (n = 8) mice during habituation and retrieval, averaged across all CS trains. Mean ± SEM (dotted line). Fold increase from 500ms pre-pip-onset. B) Average pip-induced fold increase in cross regional phase locking across the 14 CS trains during habituation and retrieval days as compared to 500 ms pre pip onset. In WT mice, an effect of Days was observed in PrL-BLA (p = 0.012) and BLA-IL (p = 0.013) while not in PrL-IL (p = 0.69). No significant effects were observed in Htr3a-KO (PrL-BLA: p = 0.223, PrL-IL: p = 0.441, BLA-IL: p = 0.691). C) Pip-induced phase locking of PrL-BLA, PrL-IL or BLA-IL theta (4-12 Hz) averaged across the first 3 trains during habituation and retrieval in WT (effect of Days: p = 0.018) and Htr3a-KO (effect of Days: p = 0.204) mice. D) Pip-evoked theta phase locking of PrL-BLA, PrL-IL, or BLA-IL in WT and Htr3a-KO during habituation day (effect of genotype: p = 0.542) or retrieval day (effect of genotype: p = 0.028). On retrieval day, Htr3a-KO showed significantly lower pip-induced PLV in BLA-IL theta as compared to WT (BLA-IL: p = 0.004). All analyses used two-way repeated measures ANOVA and subsequent Bonferroni correction when relevant. Graphs are mean ± SEM. PLV: phase locking value. *p < 0.05, **p < 0.01, ***p < 0.001.

Figure 4B displays CS-induced mean phase-locking values across the 14 successive trains (t1–t14) during habituation and retrieval in WT and *Htr3a-KO* mice, normalized to the pre-stimulus baseline. PLV values were consistently higher on the retrieval day compared to habituation in the BLA–mPFC network (PrL-BLA, BLA-IL) in WT mice, but not in *Htr3a-KO* mice. A repeated-measures ANOVA revealed a significant effect of day (retrieval vs. habituation) in WT mice, but not in *Htr3a-KO* mice, with notably higher PLV values during the first CS trains in BLA-IL interactions. Accordingly, theta phase locking averaged across the first three trains was significantly higher during retrieval than habituation in the PrL-BLA and IL-BLA networks of WT mice, but not in *Htr3a-KO* mice (Fig. 4C). While no genotype effect was observed for pip-induced theta phase locking during habituation, a significant genotype effect emerged during retrieval (Fig. 4D). During retrieval, phase locking was higher in WT mice compared to *Htr3a-KO* mice in the PrL-BLA and IL-BLA networks, reaching statistical significance only in the BLA-IL network. Altogether, these findings suggest that the CS-induced increase in theta phase locking between mPFC regions and the BLA is impaired in *Htr3a-KO* mice.

### Coupling between theta and fast-gamma in BLA during fear memory retrieval is altered in *Htr3a-KO* mice

BLA theta–gamma coupling has been shown to underlie fear learning processes in the amygdala (Stujenske et al., 2014; Cattani et al., 2024). Because theta activity in the BLA does not appear to increase during retrieval in *Htr3a-KO* mice, we next investigated whether theta–gamma coupling was affected.

In WT mice, we observed a clear increase in theta modulation of fast-gamma power on the retrieval day compared with the habituation day, with the strongest modulation occurring during the first CS presentations (Fig. 5A–B). This enhancement was weaker in *Htr3a-KO* mice, which exhibited significantly lower theta modulation of fast-gamma power in the BLA relative to WT mice during retrieval (Fig. 5C). Theta modulation of slow-gamma power did not significantly increase during pip presentations in either genotype (Fig. 5D). Nevertheless, WT mice displayed stronger theta–slow gamma coupling than *Htr3a-KO* mice on the retrieval day (Fig. 5E). Taken together, these findings indicate that the reduced theta activity observed in *Htr3a-KO* mice during retrieval is accompanied by diminished theta–gamma coupling in the BLA.

**Figure 5.**
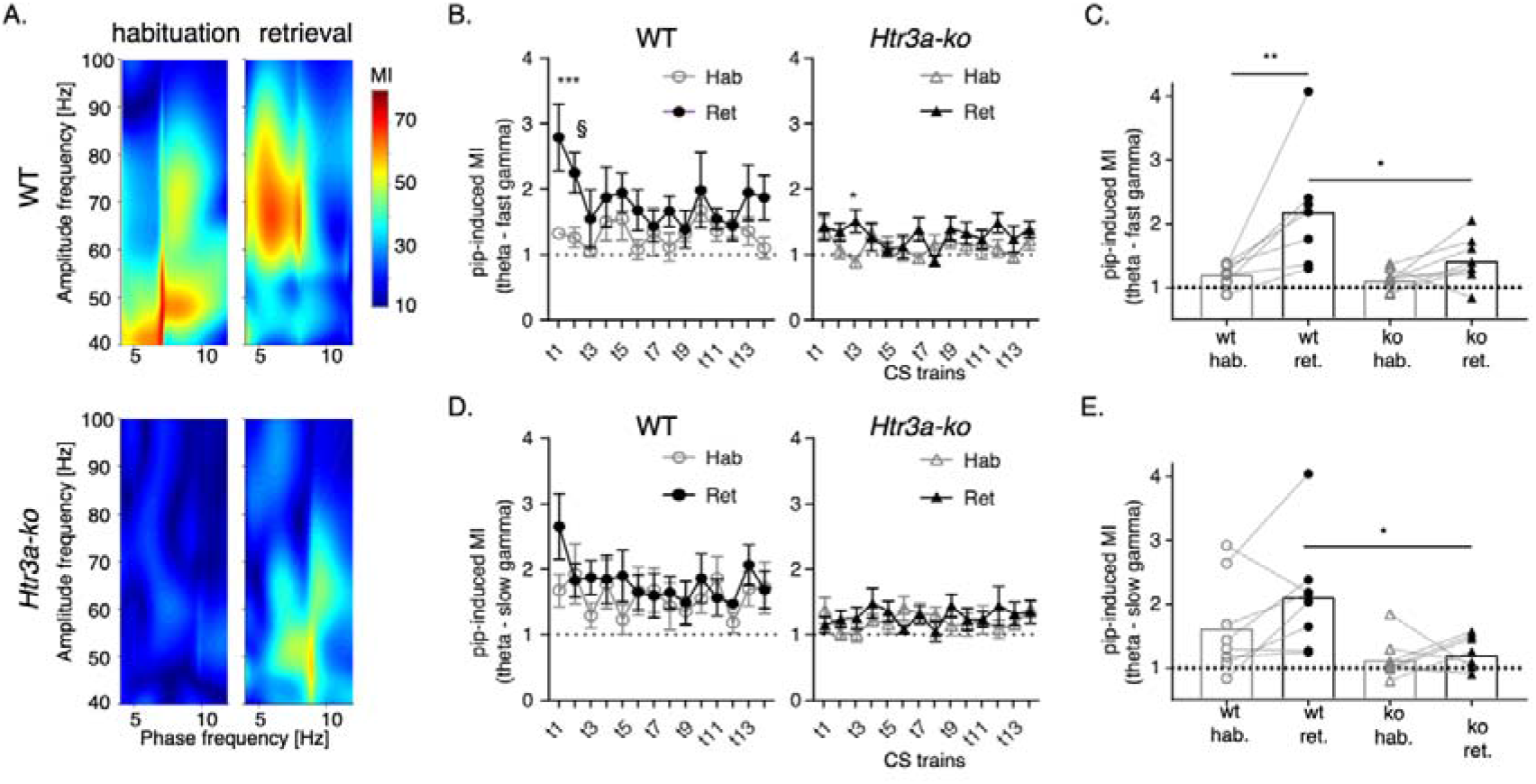
Basolateral amygdala (BLA) theta-fast-gamma coupling increases during retrieval in WT but not in Htr3a-KO. A) Phase-amplitude comodulogram of a representative BLA local field potential (LFP) recording showing theta (4-12 Hz) modulation of gamma power in WT (top) and Htr3a-KO (bottom). B) Pip-induced theta modulation of fast gamma (70-100 Hz) in the BLA across 14 CS trains during habituation and retrieval days in the WT (effect of days: p = 0.029) and Htr3a-KO (effect of days: p = 0.011) mice. Averaged fold increase from 500 ms pre pip-onset. *p < 0.05, ***p < 0.001. C) Pip-induced theta - fast gamma coupling over the first 3 trains during habituation (p>0.9) and retrieval (*p = 0.014) days in WTs (**p = 0.003) and in Htr3a-KO (p = 0.432) (main effect of days: p = 0.002; effect of genotype: p = 0.051). D) Pip-induced theta modulation of slow gamma (40-70 Hz) across 14 trains in the BLA of WT (effect of days: p = 0.261) and Htr3a-KO (effect of days: p = 0.535) mice during habituation and retrieval days. E) Average of pip-induced theta - slow gamma coupling over the first 3 trains during habituation (p = 0.135) and retrieval (*p = 0.016) days in WTs and Htr3a-KO with main effect of genotype p = 0.018. All analyses used two-way repeated measures ANOVA with Bonferroni corrections. Graphs are mean ± SEM.

## Discussion

We investigated how Htr3a receptors might contribute to the modulation of conditioned fear responses by examining behavioral adaptation and corticolimbic network dynamics during fear memory retrieval in WT and *Htr3a-KO* mice. Although *Htr3a-KO* mice acquire and express fear normally, they exhibit a marked delay in the attenuation of CS-evoked fear responses. This behavioral phenotype coincides with blunted CS-evoked activity in the PrL and IL cortices and in the BLA, including a failure to upregulate the theta power during the retrieval day. In control mice, CS-induced fear retrieval is associated with an increase in pip-evoked responses compared with the habituation period, followed by a progressive reduction across repeated stimulus trains. This difference between habituation and retrieval is absent in *Htr3a*-KO mice. In controls, the progressive decrease in stimulus-evoked activity and synchrony within the corticolimbic network likely reflects the observed attenuation of freezing during behavioral testing. Whether the absence of the initial boost in pip-evoked responses, and the consequent lack of reduction in CS-evoked responses across the stimulus trains, is causally linked to the delayed attenuation of freezing in *Htr3a-KO* mice is not causally proven here. However, it hypothesize that the altered dynamics within the fear network arise from the absence of the receptor and that this disruption contributes to the impaired adaptation of fear responses that is observed in these mice. Reduced theta phase locking across mPFC–BLA networks further supports the idea that Htr3a deletion alters functional communication between these regions. Moreover, we observed impaired theta–gamma coupling within the BLA, an oscillatory signature previously implicated in fear processing. Overall, one possible interpretation is that Htr3a receptors are necessary for enhancing theta-range activity in response to the CS and for coordinating corticolimbic synchrony during retrieval, thereby supporting efficient attenuation of conditioned fear responses.

Behaviorally, our data show that *Htr3a-KO* mice learn the fear association but exhibit a prolonged freezing response across CS presentations compared with WT animals. Previous studies have demonstrated that Htr3a receptors are not essential for fear acquisition, whether contextual or tone cued, but are required for the extinction of tone cued fear memory, as KO mice show weaker reductions in freezing during extinction sessions (Bhatnagar et al., 2004; Kondo et al., 2014). The deficit we observed in the attenuation of CS evoked freezing in Htr3a KO mice is consistent with these earlier reports of impaired extinction across retrieval sessions.

Serotonin is thought to regulate synaptic plasticity, spike synchrony, and theta oscillations in the BLA through actions on different subcellular compartments of principal neurons and distinct GABAergic interneuron populations (Bocchio et al., 2016). Htr3a receptors are expressed in a substantial population of interneurons in both the BLA and the PFC (Mascagni and McDonald, 2007; Puig and Gulledge, 2011). The reduced CS evoked LFP responses in *Htr3a-KO* mice indicate altered firing of local neurons, possibly resulting from the role of HTR3A-positive interneurons in both inhibition and disinhibition of local circuits (Tremblay, Lee et al. 2016). To answer these issues, experiments in which Htr3a receptors are specifically deleted in a subpopulation of inhibitory neurons will need to be conducted in the future.

The mPFC and BLA are anatomically and reciprocally connected, and this circuit has been strongly implicated in the modulation of fear responses (Adhikari et al., 2015; Manoocheri and Carter, 2022). While anatomical connectivity enables long-range interactions, theta synchrony allows modulation on behavioral timescales and increases within the mPFC–BLA circuit during fear states (Seidenbecher, Laxmi et al. 2003, Lesting, Narayanan et al. 2011, Likhtik, Stujenske et al. 2013, Stujenske, Likhtik et al. 2014, Karalis, Dejean et al. 2016). Our finding that WT mice exhibit increased mPFC–BLA theta coherence during the first three CS presentations on the retrieval day, compared with habituation, aligns with previous work showing increased pip-induced theta coherence during fear memory retrieval in animals that discriminate aversive from non-aversive cues (Stujenske, Likhtik et al. 2014).

Previous studies demonstrated that IL sends monosynaptic inhibitory projections to the BLA and regulates the attenuation of recent fear memories, whereas PrL sends excitatory projections that facilitate fear expression (Herry et al., 2008; Maren et al., 2013; Awad et al., 2015). The balance of activity between BLA–PrL and BLA–IL pathways determines the relative expression or attenuation of fear responses during extinction retrieval (Senn et al., 2014). Thus, the deficit in PrL–BLA synchrony in *Htr3a*-KO mice may reflect the role of Htr3a receptors in controlling fear expression, serving as a neural correlate of the heightened fear response in these animals. Although our study did not directly examine extinction learning, the reduced theta synchrony in the BLA–IL network could relate to the impaired attenuation of CS evoked responses in *Htr3a*-KO mice (Kondo et al., 2014). We also observed diminished theta–gamma coupling in the BLA during retrieval, potentially reflecting impaired feedback and feedforward processing within amygdalo–cortical circuits. Indeed, transitions between fear and safety states are mediated by shifts in BLA gamma coupling to competing theta frequency inputs (Stujenske et al., 2014).

### Limitations of the study

Electrophysiological recordings were performed under head fixed conditions rather than in freely moving animals. Although this is a limitation, this approach provided high quality signals free from movement related artifacts. The method has been used successfully in previous studies and revealed clear responses to the conditioned stimulus that are indicative of fear related neural activity. For example, a subpopulation of HTR3A- and NDNF-positive interneurons in layer 1 of auditory cortex respond to CS–US associations during discriminative fear memory retrieval in head fixed conditions (Poorthuis et al., 2018). Their activity corresponds to increased pupil dilation, a validated behavioral readout of fear in humans during CS presentation (Leuchs et al., 2017). In our experiments, we observed an increase in pupil radius during the first three CS presentations during fear memory retrieval, which then decreased to habituation levels. Although this increase reached statistical significance only in *Htr3a-KO* mice, these observations support the validity of pupil dynamics as a behavioral measure in our paradigm.

## Supplementary figure

**Supplementary Figure 1.**
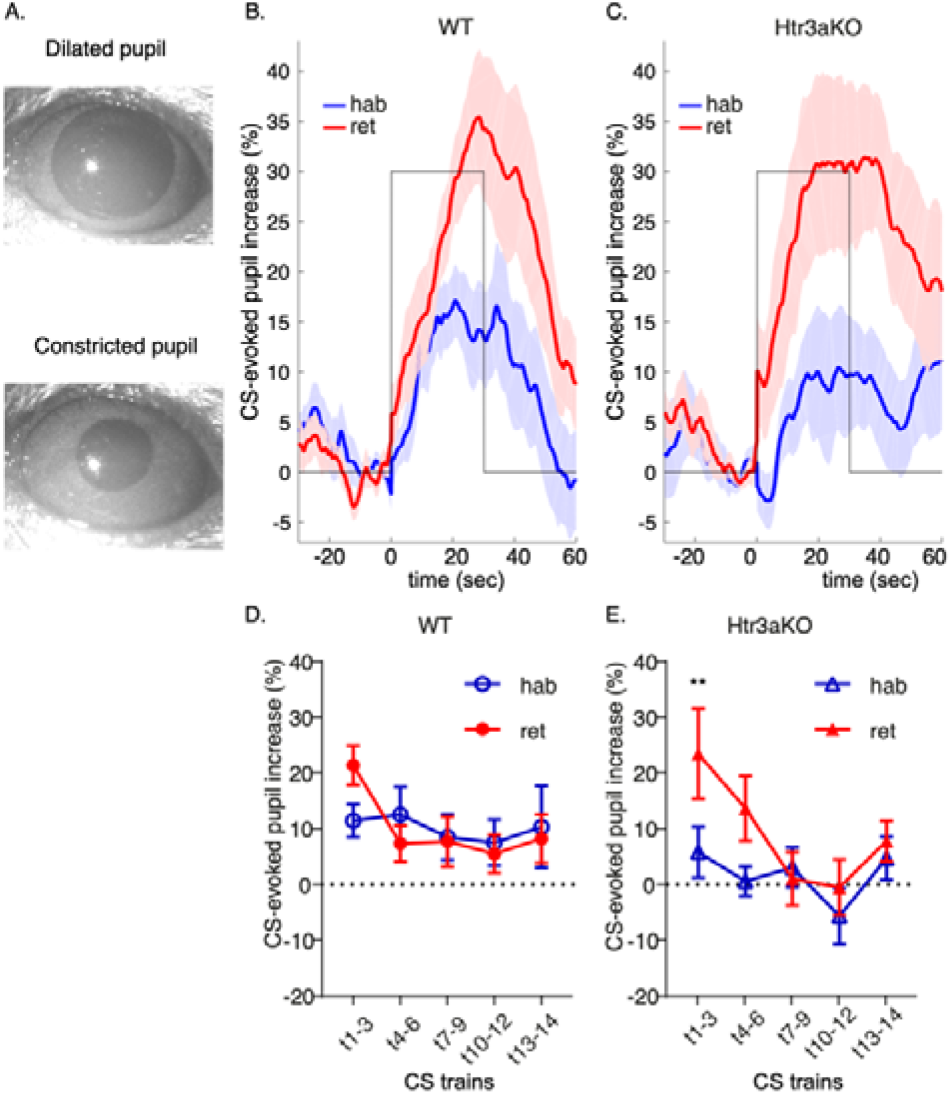
Conditional Stimulus (CS)-evoked pupil responses. To ensure that CS-fear responses could be induced in the head-fixed experimental setting, we measured pupil dilation as a CS-induced behavioral readout. Associative learning between the CS (train of pips) and the US produces reliable behavioral outputs, such as innate defensive motor behaviors (e.g., freezing), and is accompanied by signs of autonomic nervous system activation, including pupil dilation (LeDoux 2000). A) Representative examples of pupil response during CS presentation (dilated pupil) and outside of CS presentation (constricted pupil). B-C) Average of CS-evoked pupil response during the first 3 CS presentations (trains t1-t3) on habituation (blue) and retrieval (red) days in WT (n = 12) and *Htr3a-KO* (n = 8) mice. Pupil size is shown as a percentage increase from 10secs pre-CS onset. D-E) Mean CS-evoked pupil increase per block of CS presentations. Strong increases in pupil size were observed during both habituation and retrieval days, mainly for the first 3 presentations (t1-t3), although this increase reached significance only in the *Htr3a-KO* (retrieval day, WT: p = 0.071, *Htr3a-KO*: **p < 0.01; Two-way repeated measures ANOVA with Bonferonni corrections). Graphs are mean ± SEM.

